# Phosphene and Motor Transcranial Magnetic Stimulation Thresholds Are Correlated: A Meta-Analytic Investigation

**DOI:** 10.1101/2023.12.12.571304

**Authors:** P. Phylactou, T. Pham, N. Narskhani, N. Diya, D.A. Seminowicz, S.M. Schabrun

## Abstract

Transcranial magnetic stimulation (TMS) is commonly delivered at an intensity defined by the resting motor threshold (rMT), which is thought to represent cortical excitability, even if the TMS target area falls outside of the motor cortex. This approach rests on the assumption that cortical excitability, as measured through the motor cortex, represents a ‘global’ measure of excitability. Another common approach to measure cortical excitability relies on the phosphene threshold (PT), measured through the visual cortex of the brain. However, it remains unclear whether either estimate can serve as a singular measure of global cortical excitability. If PT and rMT indeed reflect global cortical excitability, they should be correlated. To test this, we systematically identified previous studies that measured PT and rMT to calculate an overall correlation between the two estimates. Our results, based on 16 effect sizes from eight studies, indicated that PT and rMT are correlated (ρ = .4), and thus one measure could potentially serve as a global cortical excitability measure. Three exploratory meta-analyses revealed that the strength of the correlation is affected by different methodologies, and that PT intensities are higher than rMT. Evidence for a PT-rMT correlation remained robust across all analyses. Further research is necessary for an in-depth understanding of how cortical excitability is reflected through TMS.

---

Since its introduction over three decades ago (Barker et al., 1985) transcranial magnetic stimulation (TMS) – a non-invasive brain stimulation technique – has been extensively utilized in research to study the nervous system (for recent reviews see Jannati et al., 2023; Sack et al., 2023). Through a coil placed over the scalp, TMS non-invasively delivers magnetic pulses, which induce changes in the ion flow of the underlaying neurons (Barker, 1991). To effectively result in neurophysiological and/or behavioral effects, TMS needs to be delivered with sufficient intensity (see Hallett, 2007; Jannati et al., 2023) that can vary across individuals depending on numerous factors that affect electromagnetic induction, such as brain-scalp distance, celebrospinal fluid thickness, electric field strength, as well as coil and/or stimulator settings (Drakaki et al., 2022; Lee et al., 2018; Wagner et al., 2009; Wang et al., 2023).

One method to determine the sufficient intensity required for a given individual is to induce TMS over the primary motor cortex (M1). TMS on M1 activates the neural circuits corresponding to the muscle represented by the targeted brain area, resulting in muscle twitching that can be measured through surface electromyography (EMG) as a motor evoked potential (MEP; Amassian et al., 1987, 1990; Barker et al., 1985). Using this methodology, TMS researchers estimate a resting motor threshold (rMT) that represents the minimum required stimulation intensity (in terms of percent of total machine output), to elicit a specified MEP amplitude in at least half the trials in the relaxed targeted muscle (typically ≥ 50μV in 5 out of 10 stimulations). In turn, the rMT serves as a proxy for measuring motor cortex excitability. For example, by using the rMT as a measure of cortical excitability, researchers can measure the effects of various interventions applied on the motor cortex, such as repetitive TMS (rTMS) protocols, to determine their excitatory or inhibitory nature (e.g., Huang et al., 2005; Traikapi et al., 2022).

A second method to determine sufficient stimulation intensity that is commonly found in the literature relates to stimulation of the occipital cortex. TMS on the primary visual cortex (V1 or V5/MT+; e.g., Pascual-Leone & Walsh, 2001) induces phosphenes^1^, which are visual percepts that appear in the visual field corresponding to the retinotopic brain area targeted by the stimulation (Walsh & Pascual-Leone, 2003). By inducing phosphenes, one can estimate a phosphene threshold (PT), which is the minimum required stimulation intensity to evoke self-reported phosphenes in at least half the trials (e.g., 5 out of 10 stimulations; see Mazzi et al., 2017). The PT has been extensively used to study vision and visual cognition (for recent reviews see Phylactou et al., 2022, 2023), as well as to measure occipital cortex excitability (for reviews see Aurora & Welch, 1998; Brigo et al., 2013).

With the exception of the use of PT for vision related research, TMS researchers most commonly rely on rMT, because it can easily produce a neurophysiological response that can be objectively measured (see Jannati et al., 2023). For example, according to safety guidelines (e.g., Rossi et al., 2009), the rMT is commonly used to define the intensity for delivering rTMS protocols, even if the targeted brain area for delivering rTMS is not within the motor cortex (e.g., De Martino et al., 2019; Traikapi et al., 2023; for a review see Lefaucheur et al., 2014). However, this approach rests on the assumption that the rMT can function as a ‘global’ measure of excitability throughout the whole brain (i.e., excitability in a different brain area is the same as that of M1, as measured through rMT). Whether this assumption can be presumed or not, remains an open question.

Previous studies have attempted to test whether the rMT reflects a global measure of cortical excitability by assessing the correlation between rMT and PT, since both measurements serve as a proxy for cortical excitability (e.g., Antal et al., 2004; Boroojerdi et al., 2002; Deblieck et al., 2008; Gerwig et al., 2003; Stewart et al., 2001; Stokes et al., 2013). Specifically, these previous studies hypothesized that if rMT and PT function as global measures of excitability, then the two measures will be correlated. The results from these previous studies were contradictory, where some studies showed a correlation between rMT and PT (Deblieck et al., 2008; Stokes et al., 2013), while others did not (Antal et al., 2004; Boroojerdi et al., 2002; Gerwig et al., 2003; Stewart et al., 2001). Given the mixed results from the previous studies, it is still unclear if rMT and PT are correlated.

The aim of this meta-analysis was to clarify whether a correlation between rMT and PT exists. To achieve this aim, relevant findings that examined the linear relationship between rMT and PT, were systematically identified, quantified, and pooled together to result in an overall correlation across all identified findings. In our pre-registration, we hypothesized that the two cortical excitability measures will not be correlated. Our findings did not support this hypothesis.

## Methods

### Study Selection

The rationale, hypothesis, and methods of the current meta-analysis were pre-registered on the open science framework (OSF; https://doi.org/10.17605/OSF.IO/TU6KH). Our meta-analysis adhered to the latest preferred reporting items for systematic reviews and meta-analyses (PRISMA) guidelines (Page et al., 2021). A systematic search of three databases (PubMed, Scopus, Web of Science) was conducted without chronological limitations and applied within the title, abstract, and keyword fields of each database. The specific search thread that was used in each database is provided in Supplementary Table 1. Additionally, the reference lists of the included studies were searched to identify any additional studies that were not captured by the database searches. No additional studies were identified through the examination of the reference lists. Moreover, seven experts in the field of TMS were contacted, in an effort to identify any potential gray literature, but no additional data were identified. Three researchers designed and completed the search strategy. Study screening and data export were evaluated by at least two independent researchers, with the utilization of Covidence (https://www.covidence.org), an online tool for evidence synthesis. Disagreements in the screening process were discussed in consensus meetings by three researchers after each phase (title/abstract screening, full-text screening, data export). In case of any remaining disagreements, a fourth researcher was to evaluate the study independently and reach a final decision, however, all disagreements were resolved during the consensus meetings.

### Inclusion and Exclusion Criteria

To be eligible for the meta-analysis, studies had to fulfill the following inclusion criteria: (1) measure both the rMT and PT from a given individual in their respective sample size, and (2) involve human participants. Studies were not included in the meta-analysis if the following exclusion criteria were met: (1) rMT and PT measured on participants with physical pathology, (2) rMT and PT measured on participants with mental and/or neurological pathology. The exclusion criteria were selected based on evidence that physical, mental, and neurological pathology can affect cortical excitability as measured by rMT (Schabrun & Hodges, 2012) and PT (Brigo et al., 2013). Studies in languages not spoken by the authors were included in the meta-analysis (i.e., Gerwig et al., 2002; Horacek et al., 2007). These studies were translated section by section using the ChatGPT 3.5 (OpenAI, 2023) large language model, using the prompt “*Translate the following text to English:*” followed by the section text in quotation marks.

### Data Analysis

The list of variables that were extracted by each study, are presented in Supplementary Table 2. Data analysis was performed using the meta package (v6.5.0; Schwarzer, 2007; see also Harrer et al., 2021) for R (v4.3.2) running on RStudio (v2023.09.1+494; R Core Team, 2023). For the meta-analysis, two researchers extracted the data independently. From each included study the reported correlation coefficient (ρ) and sample size (*n*) was extracted. Each correlation coefficient was transformed into Fisher’s *z* as shown in Equation 1:

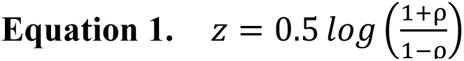

with a standard error approximated according to Equation 2:

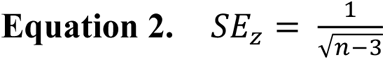

The Fisher’s *z* transformed effect sizes were used to pool an overall effect size, utilizing a random effects model (Fleiss, 1993). The overall effect size was back-transformed to ρ, for a simpler interpretation of the results. With a significance threshold set at an α < .05 level, we considered the following interpretations for the overall correlation effect size: negligible if |ρ| < .3, small if .3 ≤ |ρ| < .5, moderate if .5 |ρ| < .7, and high if .7 ≤ |ρ| (Mukaka, 2012).

Our random effects model utilized the maximum-likelihood estimator for τ^2^ and the *Q*-profile method for the τ^2^ confidence intervals. Heterogeneity was tested with the *I^2^* index, which was interpreted as low, moderate, or high, for values close to 25%, 50%, or 75%, respectively (Higgins et al., 2003). Small study bias was tested using the Egger’s test (Egger et al., 1997), accompanied by a funnel plot for visual inspection.

In addition to our registered meta-analysis, we conducted three exploratory analyses. The first exploratory analysis tested subgroup differences between studies using single pulse TMS to measure PT compared to studies using paired pulse TMS to measure PT. The second exploratory analysis investigated differences between studies estimating rMT using MEPs > 50 μV compared to MEPs > 100 μV. The differences in these two analyses were tested by comparing the estimated overall correlations with the respective χ^2^ (*df =* 1) distributions. The third exploratory analysis was conducted to explore the differences between the intensity (expressed as a percentage of maximum stimulator output) for PT and rMT. For this exploratory meta-analysis, repeated measures Cohen’s *d* (*d_rm_*) standardized differences were calculated according to Equation 3:

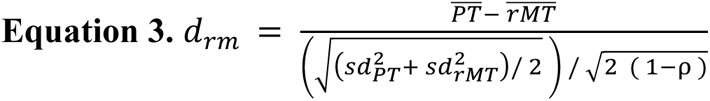

This exploratory meta-analysis was pooled using the same random-effect model described above, where positive values of the overall effect size indicate higher intensity for PT, and negative values indicate a higher intensity of rMT.

### Risk of Bias Assessment

Given that the available risk of bias (RoB) evaluation tools (e.g., Higgins et al., 2011; Sterne et al., 2016) are designed primarily to assess the quality of medical research (i.e., intervention studies), RoB was evaluated using a customized assessment tool. The customized RoB assessment tool was based on items commonly included in RoB tools (Higgins et al., 2011; Sterne et al., 2016) but also included items that are relevant to this meta-analysis specifically (i.e., TMS related; Chipchase et al., 2012). The RoB assessed 13 independent items (see Supplementary Table 2 for details), which were rated as either low bias risk, high bias risk, or unclear bias risk. RoB evaluation was completed by two researchers independently, and the final assessment scores were decided through a consensus meeting. To estimate the overall risk of bias, we calculated the sum for each study (where low risk = 0, unclear risk = 1, high risk = 2). The six sub-items of “Replicability” together with “Neuronavigation” were pooled together into one item (referred to as “Replicability”) using the same approach. For each study, the overall score from our tool was rated as low risk (Σ_RoB_ < 5), unclear risk (5:≤ Σ_RoB_ < 10), or high risk (10:≤ Σ_RoB_).

### Deviations from Registration

In our pre-registration, we reported that equivalence testing (see Lakens et al., 2020) was to be conducted if the overall effect size failed to reach significance. Accordingly, the null hypothesis was to be accepted if the 90% confidence intervals of the overall effect size included the smallest correlation of interest within their bounds (-0.3 and 0.3). However, equivalence testing was not required since all analyses reached our significance threshold of α < .05. Further, we planned to examine small study bias using the Duval-Tweedie trim and fill method (Duval & Tweedie, 2000), however this was not necessary as small study bias was not found.

### Data Availability Statement

All data and data analysis scripts used for this meta-analysis are openly available through https://osf.io/s5cpf/.

## Results

The database and reference search were completed on 10 October 2023. Following the removal of 273 duplicate reports, 574 articles were screened in the title and abstract level, out of which 559 were excluded. Fifteen full texts were assessed for eligibility, out of which five were excluded because there were no measures of PT (*k* = 2; Gamboa Arana et al., 2020; Klöppel et al., 2015) or rMT (*k* = 3; Cengiz et al., 2022; Salminen-Vaparanta et al., 2014; Sparing et al., 2005) in healthy populations. Two studies (Afra et al., 1998; Khedr et al., 2006) that met our inclusion criteria had to be excluded because the data were not provided. The corresponding authors of these two studies were contacted via email up to two times within four weeks, with a request for the data, but the data could not be retrieved. The identification and screening phases are presented as a PRISMA flowchart in Figure 1. A summary of the included studies is presented next.

**Figure 1.**
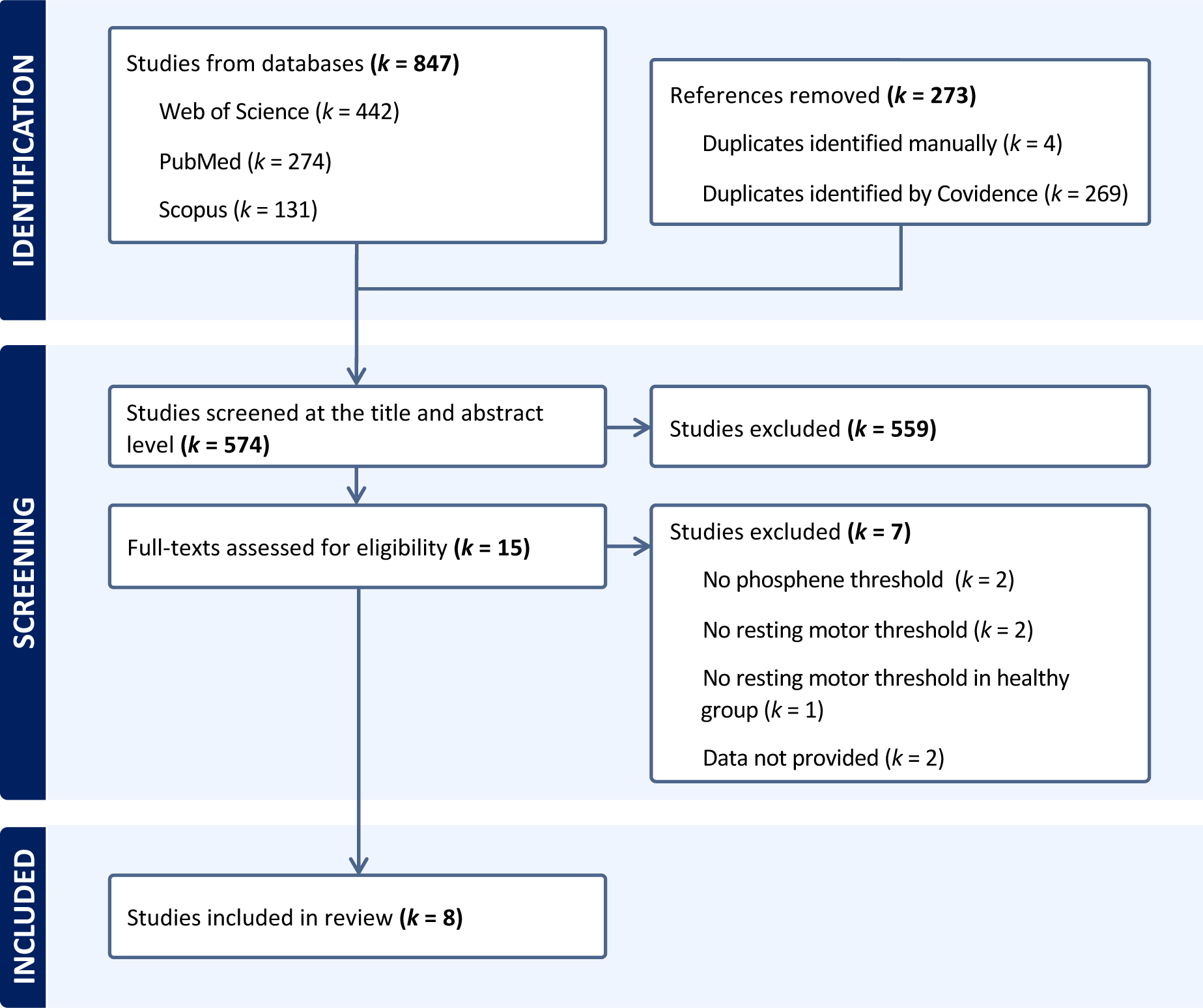
The PRISMA flowchart illustrating the identification of the eight studies that were included in the meta-analysis.

### Study Characteristics

The methodological details for each identified study are presented in Table 1. The eight identified studies included a total of 152 participants, with an average participant age of 28 years (*sd* = 2.1; from *k* = 7; see Table 1). The studies used a variety of TMS apparatuses and the use of a neuronavigation system was only reported in two studies (Horacek et al., 2007; Stokes et al., 2013).

**Table 1.**
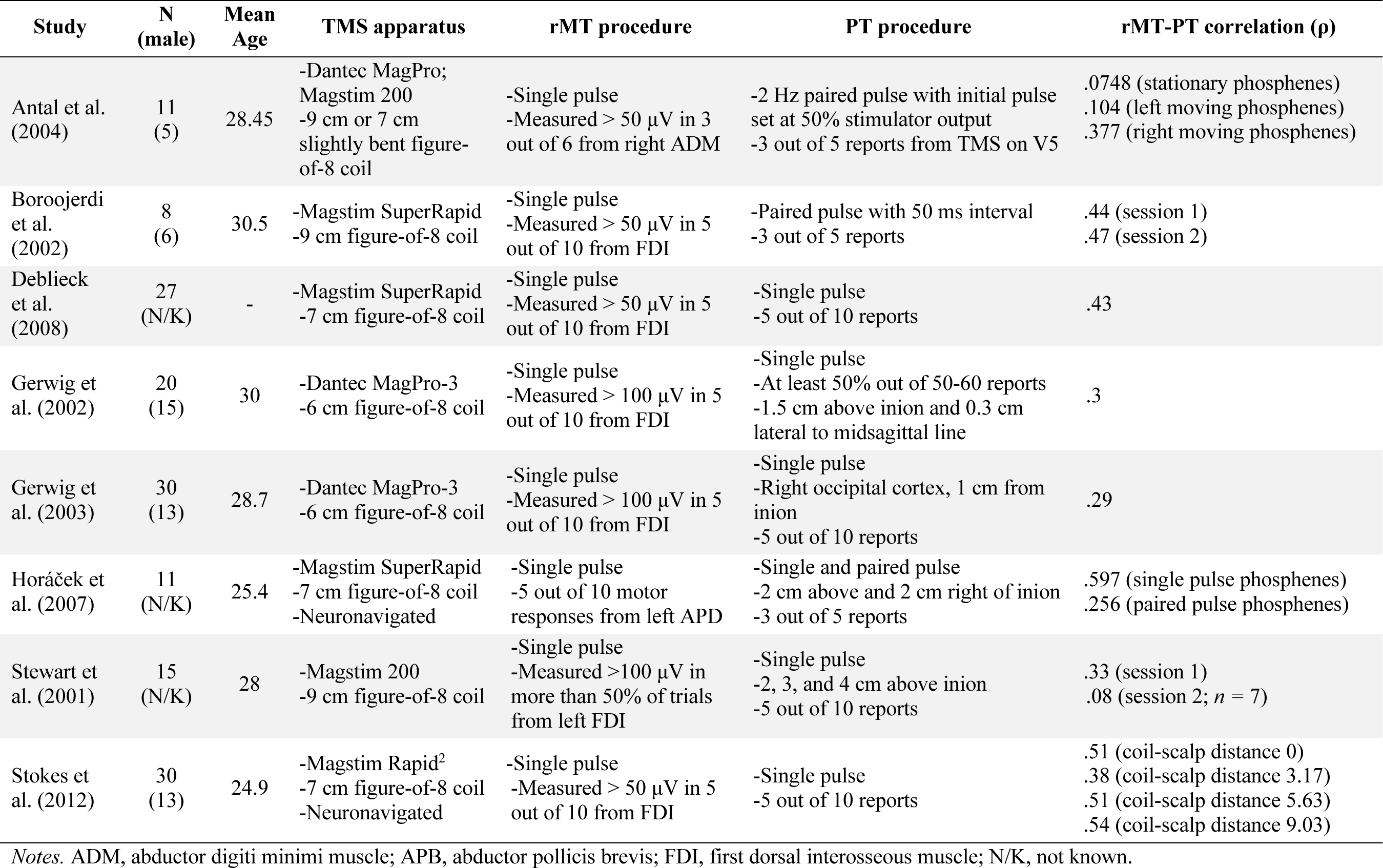
Characteristics of the studies included in the meta-analysis.

In all studies, rMT was estimated using single pulse TMS from the first dorsal interosseous (FDI; Boroojerdi et al., 2002; Deblieck et al., 2008; Gerwig et al., 2002, 2003; Stewart et al., 2001; Stokes et al., 2013), abductor digiti minimi (ADM; Antal et al., 2004), or abductor pollicis brevis (APB; Horacek et al., 2007) muscles. Four studies calculated rMT as the intensity eliciting MEPs > 50 μV in at least 50% of trials (Antal et al., 2004; Boroojerdi et al., 2002; Deblieck et al., 2008; Stokes et al., 2013), while three studies calculated rMT based on MEPs > 100 μV in at least 50% of trials (Gerwig et al., 2002, 2003; Stewart et al., 2001), and one study calculated rMT as the intensity that produced visual motor responses in at least 50% of trials (Horacek et al., 2007).

Most studies used single pulses to generate phosphenes. Two studies used paired pulses (Antal et al., 2004; Boroojerdi et al., 2002), while one study used both single pulse and paired pulse TMS (Horacek et al., 2007). One study targeted area V5 to generate moving phosphenes (Antal et al., 2004), while the remaining targeted above and lateral to the inion to generate stationary phosphenes. Next, we discuss the results of our meta-analysis that estimated an overall correlation for PT and rMT based on these studies.

### Meta-Correlation Analysis

The eight primary studies resulted in 16 effect sizes, which were pooled using a random-effects model (Figure 2A). The meta-analysis indicated that PT and rMT are correlated, ρ = .40, 95% CI = [.33, .47], *p* < .0001, which according to our registered interpretation corresponds to a small correlation (Mukaka, 2012). As reflected by the Drapery plot, this effect was robust with high confidence, beyond the α = .01 level (Figure 2B). Additionally, no evidence for heterogeneity was found, as reflected by *I^2^ =* 0% and τ^2^ = 0, *p* = .98. The estimated prediction interval indicated that future studies are likely to result in small to moderate correlations (*Prediction Interval* = [0.28, 0.51]). A visual inspection of the funnel plot (Figure 2C), as well as the results of the Egger’s test (*t_14_* = -1.67, *p* = .12), indicated no violation of symmetry, and thus there was no evidence for small study bias. Taken together, the results of our meta-analysis indicate with high confidence that PT and rMT are correlated. We further explored the relationship between PT and rMT in three exploratory meta-analyses, which we describe next.

**Figure 2.**
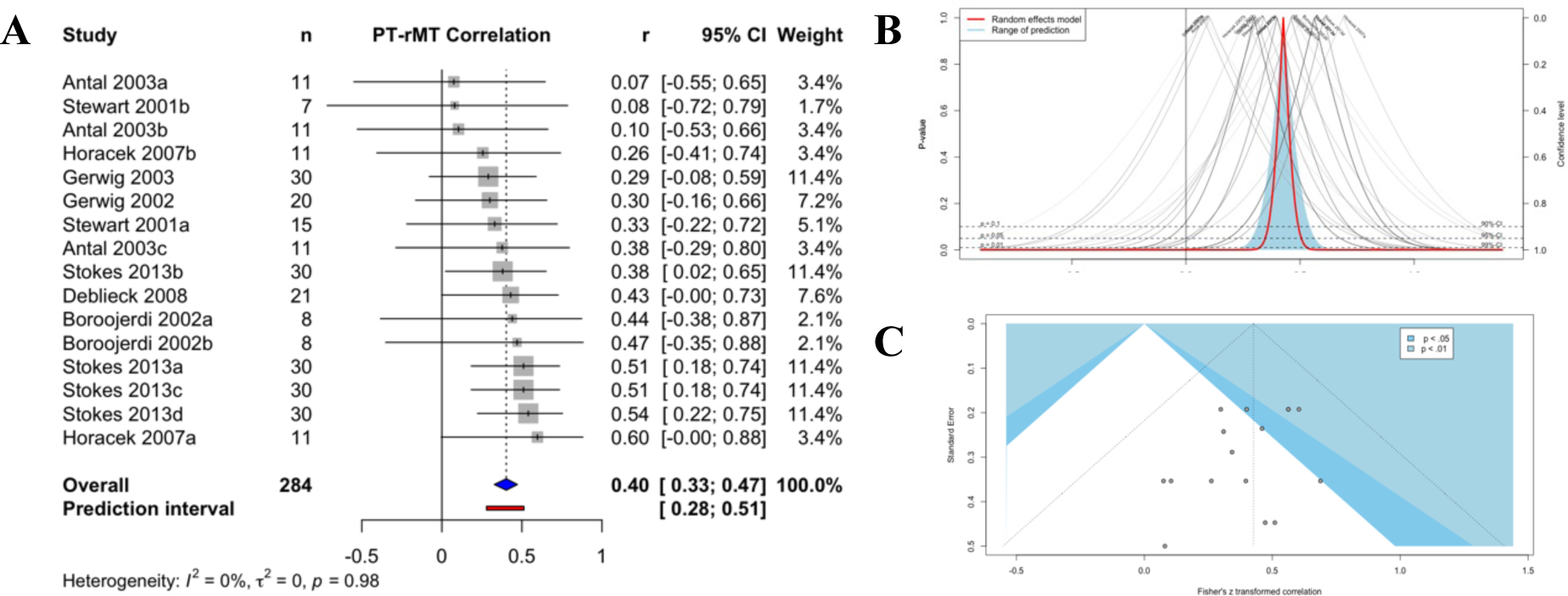
Results of the meta-analytic correlation. **(A)** The random effect model from 16 effect sizes calculated an overall effect size of ρ = .40, *p <* .0001. No heterogeneity was found (*I^2^* = 0, *p =* .98). **(B)** The Drapery plot of the overall correlation (red line) and prediction interval (blue filled line) indicate a robust effect, by plateauing beyond the α = .01 level. **(C)** A symmetrical funnel plot indicated no small study bias (*t_14_* = -1.67, *p* = .12).

### Exploratory Analyses

Three exploratory meta-analyses were conducted. The first exploratory meta-analysis investigated the differences between studies inducing phosphenes with single pulses compared to paired pulses. Our findings indicated a difference in the correlations between paired-pulse PT and single pulse PT (χ^2^_1_ = 4.48, *p* = .03), with the correlations between paired-pulse PT and rMT being smaller (ρ = .27, 95% CI = [.09, .44]; Figure 3A top) compared to single pulse PT and rMT (ρ = .43, 95% CI = [.34, .51]; Figure 3A bottom). Similarly, our second meta-analysis tested the differences between studies estimating rMT based on MEPs > 50 μV compared to MEPs > 100 μV. This analysis excluded the study by Horacek and colleageus (2007), which did not employ EMG. The results confirmed a difference between the two methods of calculating rMT (χ^2^_1_ = 8.48, *p* < .01), where rMTs calculated based on MEPs > 50 μV were more strongly correlated to PTs (ρ = .44, 95% CI = [.35, .52]; Figure 3B top) compared to MEPs > 100 μV (ρ = .29, 95% CI = [.18, .39]; Figure 3B bottom). These findings showcase that threshold estimates can vary depending on various TMS and EMG parameters.

**Figure 3.**
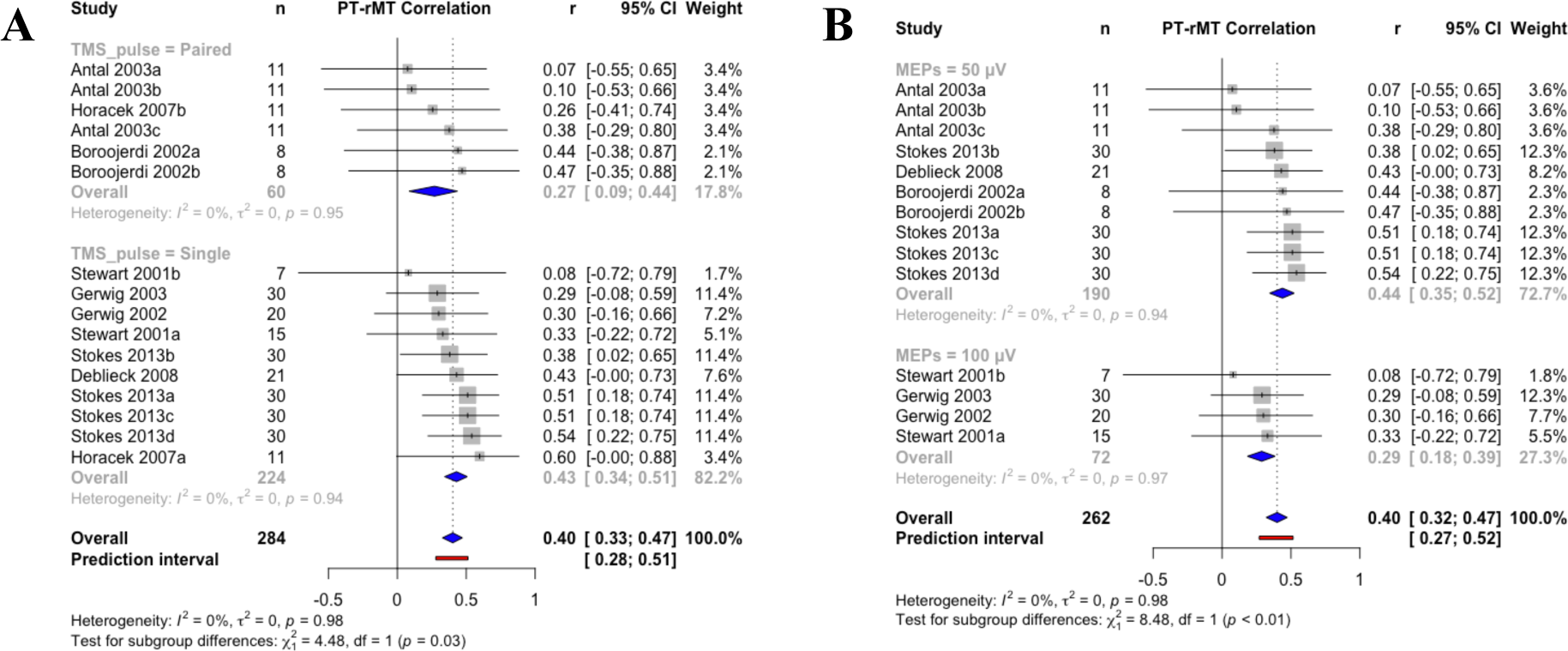
Results of the exploratory meta-analyses investigating correlation differences for different parameters of phosphene threshold and motor threshold determination. **(A)** Phosphene threshold estimated using paired pulses show a smaller correlation with resting motor threshold (ρ = .27; top) compared to single pulse phosphene threshold (ρ = .43; bottom), χ^2^_1_ = 4.48, *p =* .03. **(B)** Resting motor threshold estimated using motor evoked potentials > 50 μV show a higher correlation with phosphene threshold (ρ = .44; top) compared to > 100 μV (ρ = .29; bottom), χ^2^_1_ = 8.48, *p <* .01.

The third exploratory meta-analysis tested the differences between the PT and rMT intensities. This analysis included seven effect sizes, derived from a subset of studies (*k* = 6) that provided the required data to calculate a standardized difference coefficient^2^. A large effect size provided evidence of higher PT compared to rMT, *d_rm_* = 1.05, 95% CI = [0.06, 2.04], *p* = .0399 (Figure 4A). The Drapery plot confirmed a robust effect (Figure 4B). Notably, large heterogeneity was found for this exploratory meta-analysis, *I^2^ =* 88%, 95% CI = [79%, 93%], *p* < .01. This heterogeneity was driven by one study (Horacek et al., 2007), which was the only study showing a reversed effect (rMT > PT). Removing this study from the meta-analysis reduced heterogeneity (*I^2^ =* 12%, 95% CI = [0%, 74%], p = .335) and revealed a large effect in favor of a difference (*d_rm_* = 1.63, 95% CI = [1.22, 2.04], *p* < 0001). These results indicate that different intensities are required to elicit different neurophysiological effects. Due to the small number of identified studies, no further exploratory analyses could be conducted. Next, we present the results of the RoB assessment.

**Figure 4.**
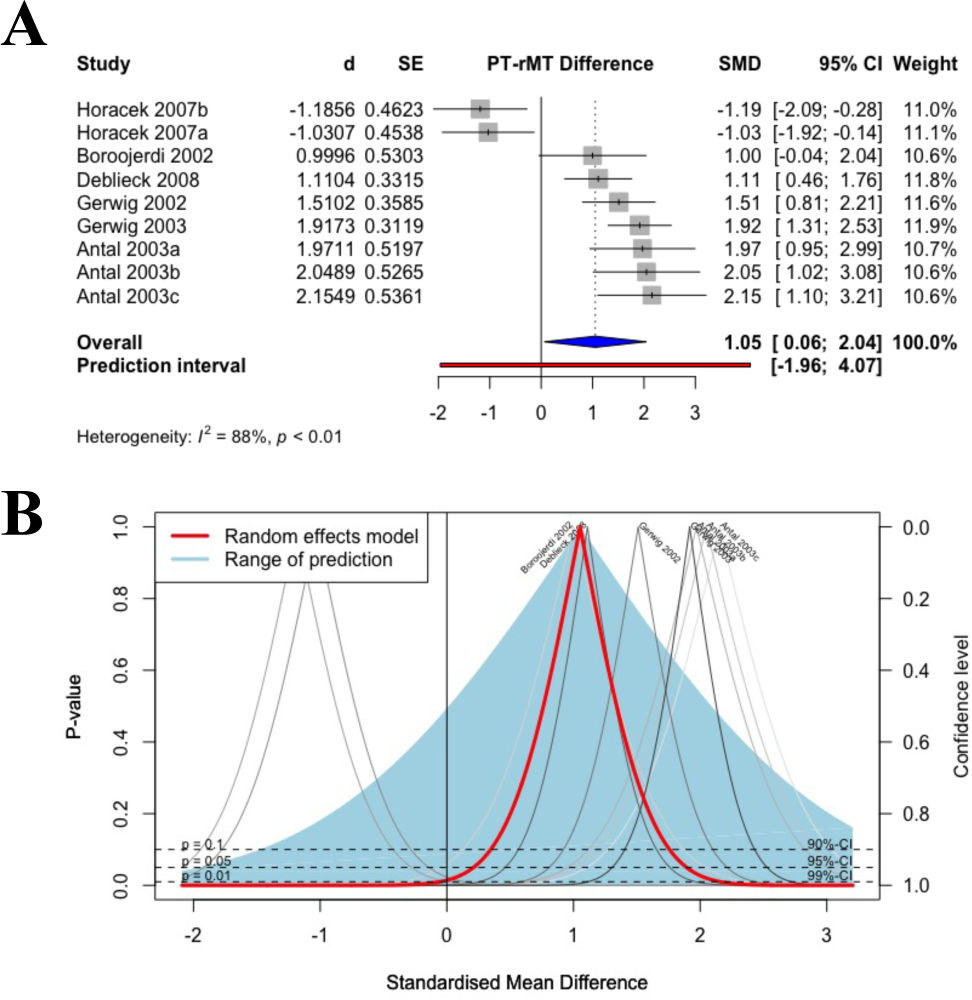
Results of the exploratory meta-analysis investigating the differences between phosphene threshold and resting motor threshold intensities. **(A)** The random effect model from 9 effect sizes provided evidence that phosphene threshold intensities are higher compared to resting motor threshold intensities *d_rm_* = 1.06, *p =* .0399. **(B)** The Drapery plot of the overall difference (red line) indicate a robust effect, by plateauing beyond the α = .05 level, however the prediction interval (blue filled line) suggests that future studies may not show this difference.

### Risk of Bias

The quality assessment indicated that all the included studies were susceptible to some degree of bias (Figures 5A and 5B). No study was ranked with overall low risk of bias. In detail, no study was registered (with the exemption of one study, which registration status was unclear; Stewart et al., 2001), and, in all studies, sample sizes were insufficient or unjustified. Further, only two studies provided sufficient detail to enable replication (Antal et al., 2004; Stewart et al., 2001), and only two studies clearly disclosed conflict of interest (Boroojerdi et al., 2002; Gerwig et al., 2003). Overall, the identified bias indicates that the quality of the included studies is not satisfactory. We next turn to a discussion of our findings.

**Figure 5.**
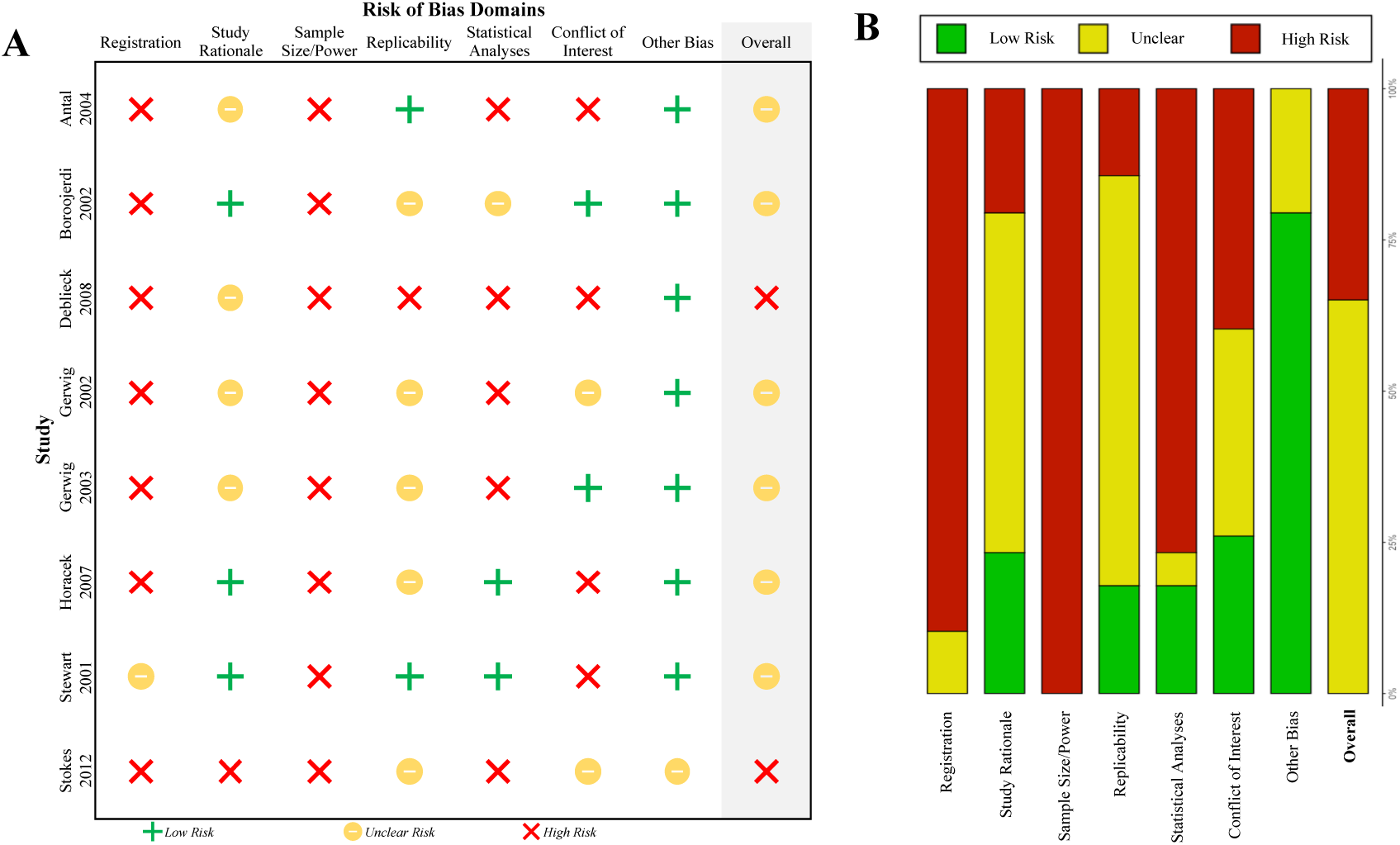
Risk of bias assessment for the included studies. **(A)** Traffic light plot indicating the assessment of each individual study and **(B)** summary plot indicating the identified bias across items for all studies.

## Discussion

After systematically identifying studies that measured the correlation between PT and rMT, we conducted a meta-analytic correlation and showed that PT and rMT are correlated. We further showed that PTs estimated with single pulses are more strongly correlated with rMT, compared to PTs generated with paired pulses and that rMTs calculated based on MEPs > 50 μV are more strongly correlated with PT, compared to MEPs > 100 μV. Moreover, our findings suggest that intensities of PT (in terms of percentage of maximum stimulator output) are higher compared to intensities of rMT. These findings, provide insight for the field of TMS research.

The correlation between PT and rMT indicates that, at least partly, global cortical excitability can be inferred through measurements of specific brain areas such as the motor or visual cortex. This provides empirical evidence in support of the approach of relying on rMT as an objective proxy of cortical excitability, even when the target area which will receive TMS (and/or rTMS) falls outside the motor region of the brain (for examples see Lefaucheur et al., 2014; see also Rossi et al., 2009). However, and as we elaborate on potential caveats below, this interpretation needs to be considered with care.

Our first exploratory meta-analysis indicated that different pulse parameters can likely result in different threshold estimates, which in turn can affect the linear relationship between PT and rMT. Similarly, our second exploratory meta-analysis revealed that different MEP determinants can affect the strength of the correlation between PT and rMT. These results are in line with a plethora of previous findings indicating how various parameters such as current direction, pulse duration, pulse wavelength, environmental conditions (e.g., light vs. dark), and threshold algorithms can affect the estimation of PT (de Graaf et al., 2017; Kammer, Beck, Erb, et al., 2001; Kammer et al., 2005; Kammer & Baumann, 2010; Mazzi et al., 2017; Ray et al., 1998) and rMT (Julkunen et al., 2009; Kammer, Beck, Thielscher, et al., 2001; Kantor et al., 1994; Peterchev et al., 2013; Tranulis et al., 2006; B. Wang et al., 2023). As such, to understand how one cortical excitability measurement may be used as a global brain proxy, an in-depth understanding of how different parameters affect the relationship between PT and rMT is necessary. Future research should aim to unravel how this relationship is affected by different parameters. As an example, future studies could explore which parameters result in approximately the same estimate for PT and rMT.

As shown in our third exploratory meta-analysis, even though the two measures of excitability share a linear relationship, rMTs and PTs are quantitatively different. Indeed, even with a perfect linear relationship (i.e., ρ = 1), a correlational analysis does not provide any information in regards to the quantitative differences between the two measures (Phylactou, Papadatou-Pastou, et al., 2022). Here, we showed that PTs were estimated as higher intensities in comparison to rMTs. This could be attributed to numerous explanations.

One potential explanation relies on the subjectivity of the threshold determination procedures. Specifically, during rMT determination, the goal is often to measure a minimal MEP (Spampinato et al., 2023), which is captured by the EMG with high precision. Despite the fact that MEPs are considered an objective measure (Jannati et al., 2023), their objectivity is violated due to some limitations. For example, the choice of the minimum peak-to-peak amplitude (e.g., 50 μV, 100 μV) for determining rMT is arbitrary, and the MEPs are prone to high trial-by-trial variability (Darling et al., 2006; Goetz et al., 2014; for a recent review see Spampinato et al., 2023). In the case of PT, researchers rely on participants’ self-reported responses, which due to subjectivity, may require a stronger neurophysiological response to elicit a positive response (Kammer et al., 2005). In other words, people may fail to report the “minimal” phosphene if uncertain. This uncertainty is also echoed by the estimation that one in four people will fail to report any reliable phosphenes (Phylactou et al., 2023).

Another explanation of this difference may lay in the neuroanatomy of the motor and visual systems. In detail, motor areas controlling the muscles commonly targeted by TMS (e.g., FDI) are close to the scalp (e.g., Gordon et al., 2023) and thus might require lower intensities to produce a neurophysiological response, whereas the primary visual cortex (i.e., V1) which extends deeper into the brain through the calcarine sulcus, may require higher intensities to produce a noticeable response. The above potential explanations could be explored by future work, by, for example, comparing input-output curves for MEPs and phosphene characteristics (for example see Pascual-Leone & Walsh, 2001).

It is also possible that motor and visual areas have different excitability thresholds. For example, even though brain plasticity is presumed to be globally controlled through GABAergic processes in the visual (Greifzu et al., 2014; Harauzov et al., 2010) and motor (McLean et al., 1996; Schabrun & Hodges, 2012; Thapa et al., 2021; Ziemann et al., 2001) system, it has been shown that when plasticity is elicited simultaneously for the visual and motor system, motor plasticity may remain intact at the cost of visual plasticity (Sarı & Lunghi, 2023). However, the consistent findings of a correlation between PT and rMT across all our analyses, indicate that, even if qualitatively different, the two cortical excitability measures are indeed described by a linear relationship.

It should also be mentioned that even though the correlation between PT and rMT suggest that global cortical excitability may be reflected by the either of the two cortical excitability measures, our exploratory analyses showcase the complexity of such linearity. Given the lack of our current understanding of how one estimate can precisely reflect another, it is advised that researchers adjust their parameters for estimating thresholds according to their specific aims in a given study (Jannati et al., 2023). Online tools, such as TMS-SMART (Meteyard & Holmes, 2018), and simulation tools such as sim-NIBS (Saturnino et al., 2019), can help researchers make more informed decisions regarding their TMS parameters. Further, closed looped TMS designs (e.g., through concurrent electroencephalography or functional magnetic resonance imaging) may enhance the reliability and effectiveness of TMS and rTMS procedures (Peters et al., 2020; Sack et al., 2023).

Attention should also be brought to the identified RoB of the primary studies included in this review. Our quality assessment revealed that all studies were prone to at least some degree of bias risk. This finding echoes the need for the scientific community to adhere to specific TMS reporting criteria (Chipchase et al., 2012), to enable replicability, minimize bias, and facilitate TMS protocol standardization. Though, it should also be noted that our RoB assessment may also be susceptible to bias. In detail, the current widely used RoB assessment tools are tailored to appraise mainly clinical related research (e.g., Higgins et al., 2011; Sterne et al., 2016) and relevant RoB tools for basic experimental research are very limited. As such, we had to develop our own RoB tool, which was adjusted by previously developed tools (Chipchase et al., 2012; Higgins et al., 2011; Sterne et al., 2016). In the future, meta-scientific work could aim to develop RoB and other quality assessment tools, which will be utilized to evaluate basic experimental work. This is timely, considering the potential uprise of meta-analytic methods for basic science questions (see Mikolajewicz & Komarova, 2019; Phylactou, Traikapi, et al., 2022).

Collectively our findings provide support for the idea that rMT and PT may serve as a global measure of excitability. However, these findings need to be considered in light of our study’s limitations. Firstly, correlation analysis itself does not enable for strong conclusions or inferences to be drawn. As mentioned above, even when two variables are correlated, they can still be quantitively very different (Phylactou, Papadatou-Pastou, et al., 2022). This was also reflected in our exploratory meta-analyses that illustrated how the strength of the correlation changed when considering different thresholding parameters, and by showing that PT was overall higher than rMT. Further, it has been shown that signals from slowly evolving systems (such as cortical excitability) can potentially lead to false correlations (Meijer, 2021). Despite the limitations of correlation analysis, our finding was robust across all analyses, thus confirming that, at least in part, PT and rMT are correlated.

Secondly, our findings are limited to the healthy brain. The choice of excluding studies focusing on the damaged brain was based on findings that physical, mental, and/or neurological pathology can affect cortical excitability as revealed by TMS measures (e.g., Brigo et al., 2013; Schabrun & Hodges, 2012). However, the question of whether rMT and PT can be used as global measures of cortical excitability is as important for clinically relevant applications, considering the wide use of TMS in applied clinical work (see Lefaucheur et al., 2014; Sack et al., 2023). Given that this was, to the best of our knowledge, the first attempt to systematically identify and analyze studies that measured the relationship of PT and rMT, our meta-analysis serves as a foundation for further exploration. We suggest that future meta-analyses build on our findings and attempt to understand the relationship of PT and rMT further. For example, since our findings indicate the presence of a correlation between PT and rMT, subsequent studies can investigate whether a similar relationship exists in the damaged brain.

## Conclusions

In conclusion, in one meta-correlational analysis and three exploratory analyses we showed that PT and rMT are correlated. Even though the strength of this linear relationship is affected by different TMS parameters, the correlation remained, showcasing the robustness of this relationship. Our findings suggest that one TMS measure, such as PT or rMT, may be used as a global measure of cortical excitability. However, future work is required to further understand the complexity of cortical excitability and the different TMS measures.

## Author Contributions

PP: conceptualization, data curation, formal analysis, methodology, visualization, funding acquisition, writing original draft; PT: data curation, analysis support; NN: data curation; ND: data curation; DAS: conceptualization, methodology, analysis support and review, writing review and editing; SMS: conceptualization, methodology, analysis support and review, funding acquisition, writing review and editing.

## Conflict of Interest

The authors have no conflict of interest to declare.

References marked with “*” were included in the meta-analysis.

## Footnotes

1 It has also been demonstrated that phosphenes can be induced from stimulating higher order brain areas such as the posterior parietal cortex (Fried et al., 2011).

2 Data for Horacek and colleagues (2007) were estimated from medians and interquartile ranges, based on approximations for means (Luo et al., 2018) and standard deviations (Wan et al., 2014).

