## Supplementary Materials for "Phosphene and Motor Transcranial Magnetic Stimulation Thresholds Are Correlated: A Meta-Analytic Investigation"

**Supplementary Material**

v1, 12 Dec 2023

**Supplementary Table 1:** Search threads used in each database for study identification.

| | |
|---|---|
| Web of Science | (((((((TS=(transcranial magnetic stimulation)) OR TS=(tms)) AND TS=(threshold)) OR TS=(excit*)) AND TS=(motor)) AND TS=(visual)) OR TS=(phosphen*)) AND TS=(correl*) |
| Scopus | TITLE-ABS-KEY ( transcranial AND magnetic AND stimulation OR tms ) AND TITLE-ABS-KEY ( threshold OR excit* ) AND TITLE-ABS-KEY ( motor ) AND TITLE-ABS-KEY ( visual OR phosphen* ) AND TITLE-ABS-KEY ( correl*) |
| Pubmed | ((((((('transcranial magnetic stimulation'[Title/Abstract]) OR (tms[Title/Abstract])) AND (threshold[Title/Abstract])) OR (excit*[Title/Abstract])) AND (motor[Title/Abstract])) AND (visual[Title/Abstract])) OR (phosphen*[Title/Abstract])) AND (correl*[Title/Abstract]) |

**Supplementary Table 2.** List and description of each variable that was extracted by each study.

| Sample Size (*n*) | Correlation Coefficient (*r*) | Age | Sex | TMS apparatus | TMS parameters | Neuronavigation | rMT method | PT method |
|---|---|---|---|---|---|---|---|---|
| The total sample size included for the calculation of the rMT-PT correlation. This could differ from the total study sample size (e.g., if the correlation was measured separately for a healthy control group vs. a condition group, where only the healthy group data will be used). | Raw correlation coefficient as provided in the study or as calculated by the authors of the meta-analysis by the provided data. | Mean age as reported in the primary study or as calculated by the authors of the meta-analysis by the provided data. If a range is provided in a primary study, then the midpoint of the range will be used as the approximate mean age of that primary study. | The reported sex (male/female/other) from each primary study as reported in the primary study or as calculated by the authors of the meta-analysis by the provided data. | The model of the TMS machine and specifications of the coil used in each primary study. | The parameters of the TMS used to calculate rMT and PT, including number of pulses, frequency (if not single-pulse TMS), and duration. | Whether or not a neuronavigation system was used for the primary study. | Description of the method used to calculate the rMT, including the targeted muscle and the MEP threshold. | Description of the method used to calculate the PT including the target area (i.e., V1 or V5). |

**Supplementary Table 3:** Risk of Bias Assessment Items.

| Assessment Item | Description |
|---|---|
| 1. Study registration. | Evaluation of whether the study's hypotheses and methods were pre-registered. Study pre-registration minimizes possible bias. |
| 2. Study rationale, aims, and hypotheses. | Evaluation of whether the study's rationale, aims, and hypotheses are stated clearly. Clear descriptions minimize possible bias. |
| 3. Sample size and power. | Evaluation of whether the sample size resulted in adequate power (frequential approach) or an adequate ratio of evidence (Bayesian or likelihoodist approach). Adequate power minimizes possible bias. |
| 4. Replicability. | Evaluation of whether adequate and clear details are provided so that the study can be replicated. Sufficient information minimizes possible bias. This item will be assessed based on the reporting of the following (adapted from Chipchase et al., 2012): (i) coil type, (ii) coil orientation, (iii) coil location and stability, (iv) stimulator type, (v) pulse shape, (vi) pulse parameters [if not single pulse]. |
| 5. Neuronavigation. | Evaluation of whether a neuronavigation system was utilized for TMS. The use of neuronavigation minimizes possible bias. |
| 6. Statistical analyses. | Evaluation of whether the statistical analyses used were appropriate (e.g., normality checks, representative priors, corrections, transformations). Appropriate analyses minimize possible bias. |
| 7. Conflict of interest. | Evaluation of whether any conflict of interest was disclosed (including funding information). Disclosing conflict of interest minimizes risk of bias. |
| 8. Other bias. | Evaluation of whether any other factor is evident that could affect bias in the study. Lack of any additional factors minimizes risk of bias. |
